## Supplementary Materials for "The interplay between the polar growth determinant DivIVA, the segregation protein ParA and their novel interaction partner PapM controls the *Mycobacterium smegmatis* cell cycle by modulation of DivIVA subcellular distribution"

**Table S1. Constructs used in the study**

| Constructs | Relevant genotype | Source |
|---|---|---|
| Constructs used in *E. coli* BTH system | | |
| pUT18C | pUC replicon, $p_{lac}$ promoter, ampicillin resistance, IPTG-inducible, T18 fragment of *cyaA* gene (encoding 225 to 399 amino acids) | (Karimova et al., 1998) |
| pKT25 | P15A replicon, $p_{lac}$ promoter, kanamycin resistance, IPTG-inducible T25 fragment of *cyaA* gene (encoding 1- 224 amino acids) | (Karimova et al., 1998) |
| pKT25*parA* | pKT25 derivative containing *M. smegmatis parA* gene | (Ginda et al., 2013) |
| pKT25*parA*T3A | pKT25 derivative containing *M. smegmatis parA*T3A gene | (Pióro et al., 2019) |
| pKT25*parA*K44A | pKT25 derivative containing *M. smegmatis parA*K44A gene | (Pióro et al., 2019) |
| pKT25*parA*D68A | pKT25 derivative containing *M. smegmatis parA*D68A gene | (Pióro et al., 2019) |
| pKT25*parA*G40V | pKT25 derivative containing *M. smegmatis parA*G40V gene | (Pióro et al., 2019) |
| pKT25*parA*R219E | pKT25 derivative containing *M. smegmatis parA*R219E gene | (Pióro et al., 2019) |
| pKT25*parB* | pKT25 derivative containing *M. smegmatis parB* gene | (Ginda et al., 2013) |
| pUT18C*divIVA* | pUT18C derivative containing *M. smegmatis divIVA* gene | (Ginda et al., 2013) |
| pUT18C*divIVA I-II* | pUT18C derivative containing a fragment of *M. smegmatis divIVA* gene used for the production of T18C-DivIVAI-II (amino acids 1-72) | (Ginda et al., 2013) |
| *pUT18CdiIVA II-III* | pUT18C derivative containing a fragment of *M. smegmatis divIVA* gene used for the production of T18C-DivIVAII - III (amino acids 1-143) | (Ginda et al., 2013) |
| pUT18C*divIVA III-IV* | pUT18C derivative containing a fragment of *M. smegmatis divIVA* gene used for the production of T18C-DivIVAIII - IV (amino acids 68-272) | (Ginda et al., 2013) |
| pUT18C*divIVA IV* | pUT18C derivative containing a fragment of *M. smegmatis divIVA* gene used for the production of T18C-DivIVAIV (amino acids 139-272) | (Ginda et al., 2013) |
| pKT25*divIVA* | pKT25 derivative containing *M. smegmatis divIVA* gene | (Ginda et al., 2013) |
| pUT18C*msmeg_5597* | pUT18C derivative containing *M. smegmatis msmeg_5597* (*papM*) gene | This study |
| pKT25*msmeg_5597* | pKT25 derivative containing *M. smegmatis* (*msmeg*_5597) *papM* gene | This study |
| Constructs used for protein purification | | |
| pGEX-6P-2 | pGEX-6P-2 ampicillin resistance, GST-tag vector | Lab stock, University of Wroclaw |

| | | |
|---|---|---|
| pGEX-6P-2*parA* | pGEX-6P-2 derivative containing *M. smegmatis parA* gene | (Jakimowicz et al., 2007) |
| pET28a | pET28a kanamycin resistance, His-tag vector | Lab stock, University of Wroclaw |
| pET28a*papM* | pET28a derivative containing *M. smegmatis papM* gene | This study |
| Constructs used for colocalization microscopy in *E. coli* | | |
| pETDuet-1 | ampicillin resistance vector, two multiple cloning sites, the pBR322-derived ColE1 replicon, *lacI* gene | Lab stock, University of Wroclaw |
| pJP108*divIVA*$_{Ms}$ | pJP108 containing *icsA507-620-mcherry* fused with *M. smegmatis divIVA* gene | (Pióro et al., 2019) |
| pETDuet-*pknB*$_{KD}$ | pETDuet-1 derivative containing fragment of *M. smegmatis pknB* gene encoding kinase domain (11-274 aa) | This study |
| pETDuet-*mcherry-divIVA* | pETDuet-1 derivative containing *icsA507-620-mcherry* fused with *M. smegmatis divIVA* gene | This study |
| pETDuet-*pknB*$_{KD}$ *mcherry-divIVA* | pETDuet-1 derivative containing fragment of *M. smegmatis pknB* gene encoding kinase domain (11-274 aa) (MCS1) and *divIVA* gene fused with *mcherry* (MCS2) | This study |
| pETDuet-*ics-mcherry* | pETDuet-1 derivative containing *icsA507-620-mcherry* | This study |
| pACYC-Duet-1 | chloramphenicol resistance vector, two multiple cloning sites, the P15A replicon, *lacI* gene | Lab stock, University of Wroclaw |
| pACYC-*egfp-parA* | pACYCDuet-1 derivative containing *egfp* gene fused with *M. smegmatis parA* gene | (Pióro et al., 2019) |
| pACYC-*papM-mT2* | pACYCDuet-1 derivative containing *M. smegmatis papM* gene fused with *mturquoise2* | This study |
| pACYC-*papM-mT2; egfp-parA* | pACYCDuet-1 derivative containing *M. smegmatis papM* gene fused with *mturquoise2* (MCS1) and *egfp* gene fused with *M. smegmatis parA* gene (MCS2) | This study |
| Constructs used for *M. smegmatis* modifications | | |
| p2Nil | kanamycin resistance, *oriE*, suicide vector for allelic replacement | (Parish and Stoker, 2000) |
| pGOAL | ampicillin resistance, *oriE*, selective PacI cassette with *lacZ*, *sacB* and *kanR* genes | (Parish and Stoker, 2000) |
| p2Nil*ΔpapM*GOAL | p2Nil derivative for deletion of *papM* gene | This study |
| p2Nil*lsr2-mT2*GOAL | p2Nil containing *mTurquoise2* gene fused with *lsr2* | (Kołodziej et al., 2021) |
| pMV306p$_{ami}$ | pMV306 containing acetamide inducible promoter p$_{ami}$ | (Ginda et al., 2013) |
| pMV306p$_{ami}$*papM* | pMV306p$_{ami}$ derivative containing *M. smegmatis papM* gene | This study |
| pMVp$_{nat}$*egfp-parA* | pMV306p$_{ami}$ derivative containing *egfp* gene fused with *parA* gene | (Ginda et al., 2013) |
| pMVp$_{nat}$PA*mcherry-parA* | pMV306p$_{ami}$ containing PA*mcherry* gene fused with *parA* under its native promoter | (Pióro et al., 2019) |

| | | |
|---|---|---|
| pKW08$p_{tet}$ | pKW08 containing tetracycline inducible promoter $p_{tet}$ | Lab stock, University of Wroclaw |
| pKW08$p_{tet}$*mcherry-divIVA* | pKW08$p_{tet}$ derivative containing *mcherry* gene fused with *divIVA* | This study |

**Table S2. Strains used in the study**

| | | |
|---|---|---|
| *E. coli* strains | | |
| DH5α | *F-, endA1, glnV44, thi-1, recA1, relA1, gyrA96, deoR,nupGΦ80dlacZΔM15Δ(lacZYA-argF) U169, hsdR17(rKmK+), λ-* | Lab stock, University of Wroclaw |
| BTH101 | F-, *cya-99, araD139, galE15, galK16, rpsL1 (Str $^r$), hsdR2, mcrA1, mcrB1* | Lab stock, University of Wroclaw |
| BL21(DE3) | *F-, ompT, gal, dcm, lon, hsdSB(rB- mB-), λ(DE3)* | Lab stock, University of Wroclaw |
| *M. smegmatis* strains | | |
| WT | *M. smegmatis* mc$^2$ 155 | lab stock, University of Wroclaw |
| KG22 | *M. smegmatis* mc$^2$ 155 *ΔparA* | (Ginda et al., 2013) |
| MP24 | *M. smegmatis* mc$^2$ 155, *attB*L5:: pMV306$p_{ami}$*ø* | (Pióro et al., 2022) |
| KG56 | *M. smegmatis* mc$^2$ 155 *ΔparA,* attBL5:: pMV306$p_{nat}$*egfp–parA* | (Ginda et al., 2013) |
| DJMP01 | *M. smegmatis* mc$^2$ 155 *ΔparA,* attBL5:: pMV306$p_{nat}$*PAmcherry–parA* | (Pióro et al., 2019) |
| IM01 | *M. smegmatis* mc$^2$ 155 *ΔpapM* | This study |
| IM02 | *M. smegmatis* mc$^2$ 155 *ΔparA, ΔpapM* | This study |
| IM11 | *M. smegmatis* mc$^2$ 155 *ΔparA,* attBL5::pMV$p_{ami}$Φ | This study |
| IM12 | *M. smegmatis* mc$^2$ attBL5::pMV$p_{ami}$*papM* | This study |
| IM13 | *M. smegmatis* mc$^2$ 155 *ΔparA,* attBL5::pMV$p_{ami}$*papM* | This study |
| IM14 | *M. smegmatis* mc$^2$ 155 *ΔparA, ΔpapM,* attBL5::pMV$p_{nat}$*egfp-parA* | This study |
| IM15 | *M. smegmatis* mc$^2$ 155 *ΔparA, ΔpapM,* attBL5:: pMV306$p_{nat}$*PAmcherry–parA* | This study |
| IM16 | *M. smegmatis* mc$^2$ attBL5::pKW08$p_{tet}$*mcherry-divIVA* | This study |
| IM17 | *M. smegmatis* mc$^2$ *ΔpapM,* attBL5::pKW08$p_{tet}$*mcherry-divIVA* | This study |
| IM18 | *M. smegmatis* mc$^2$ *ΔparA,* attBL5::pKW08$p_{tet}$*mcherry-divIVA* | This study |
| IM19 | *M. smegmatis* mc$^2$ *ΔparA, ΔpapM,* attBL5::pKW08$p_{tet}$*mcherry-divIVA* | This study |
| IMPW1 | *M. smegmatis* mc$^2$ *ΔparA, ΔpapM,* attBL5::pMV$p_{ami}$*papM* | This study |

**Table S3. Oligonucleotides used in this study**

| Name | Sequence 5' to 3' |
|---|---|
| PknB_Ms_Fw_HindIII | C**AAGCTT**gATGAGGTCGGCGCGCATC |
| PknB_Ms_Rv_EcoRI | CT**GAATTC**ATACGAACTCGGTGAGATCCTC |
| Cherry_Fw_Nde | CC**CATATG**GTGAGCAAGGGCGAGGAGG |
| Div_revSnaBI | CC**TACGTA**TCAGTTGTTGCCGCGGTTGAACTGG |
| Ics_Fw_NdeI | **CATATG**AGTACTATTCTGGCAGATAATCTCAGCCATC |
| Cherry_rvSnaBI | CC**TACGTA**TTACTTGTACAGCTCGTCCATGCCGC |
| 5597_Fw_NcoI | AACTTTAATAAGGAGATATA**CCATGG**GTGGGACGTCATCGCGAGTT |
| 5597_linker_Rv | GCCCGCCGCCGAGCCCGCCGAGCCAGGCGCTTGCGCGTGGCCTTCCCC |
| linker_mT2_Fw | AAGGCCACGCGCAAGCGCCTGGCTCGGCGGGCTCGGCGGCGGGCT |
| mt2_Rv_EcoRI | CCTGCAGGCGCGCCGAGCTC**GAATTC**TCACTTGTACAGCTCGTCCA |
| MSMEG_5597_XbaI_BamHI_Fw | CC**TCTAGA**G**GGATCC**ATGGGACGTCATCGCGAGTT |
| MSMEG_5597_KpnI_EcoRI_Rv | CC**GGTACC**CG**GAATTC**GCGCTTGCGCGTGGCCTT |
| Msmeg5597_F1_HindIII_Fw | C**AAGCTT**TATGAAATCGGCCGTTGG |
| Msmeg5597_F1_BamHI_Rv | C**GGATCC**ACGACCGGTAATAAAACTACC |
| Msmeg5597_F2_BamHI_Fw | C**GGATCC**TGGGAGTTGGGCGGTACG |
| Msmeg5597_F2_PacI_Rv | C**TTAATTAA**GCCCTGTCTGCCAGCTTG |
| 5597_pami_XbaI_Fw | CTC**TCTAGA**GTGGGACGTCATCGCGAGTTC |
| 5597_pami_KpnI_Rv | CTT**GGTACC**TCAGCGCTTGCGCGTGGC |
| RT-PCR3_Fw | ATCGCGAGTTCGATGTGGAC |
| RT-PCR3_Rv | TACCCTTTGCGCCAGAACAC |
| pKW08F2 | GGTGGTGAGTCATAGTTGC |
| pKW08RV2 | GGATTACACATGACCAACTTC |

*Boldface type indicates the restriction site.

**Supplementary Materials and Methods**

**BTH library construction, screening and interaction verification**

For BTH library construction chromosomal DNA was isolated from the *M. smegmatis* mc2 155 strain cultured on solid NB medium for 2 days. 50 µg of chromosomal DNA was sonicated in aliquots to deliver fragments with length of 500 to 1500 bp (as verified by gel electrophoresis). After sonication, aliquots of DNA were pooled and then the ends of the resulting fragments were filled with DNA T4polymerase and Klenow DNA polymerase I fragment and next phosphorylated with T4 polynucleotide kinase (PNK). After the reaction, the DNA was purified by preparative electrophoresis in a 1% agarose gel, the fragments within the size range of 600-1500 bp were excised and isolated with the QIAquick Gel Extraction Kit (Qiagen). The pKT25 vector used to construct the library was digested with the restriction enzyme SmaI (for 3 days, gradually increasing the amount of enzyme). Then DNA was precipitated and dephosphorylated and used for ligation with the obtained insert, followed by transformation of *E. coli* TOP10 competent cells. Transformants were selected on LB medium supplemented with kanamycin. Process of ligation and transformation were optimized and adjusted to ultimately obtain about 400 colonies on a single one plate. Altogether 11 ligations performed and 22 transformations delivered total of 38 000 colonies. Colonies were harvested from the obtained plates and plasmid DNA was isolated using the Qiagen Plasmid Maxi Kit (Qiagen). Finally, 1.5 ml of pKT25lib$_{MS}$ DNA at concentration 480 ng/µl was obtained.

To screen the BTH library, the electrocompetent cells of *E. coli* BTH101 strain containing pUT18*parA* were transformed with pKT25lib$_{MS}$ plasmid DNA. Transformation mixture was plated on LB medium supplemented with 0.004% X-gal, 50 µg/ml kanamycin, 100 µg/ml ampicillin, and 0.5 mM isopropyl-β-D-1-thiogalactopyranoside (IPTG). The dilutions of the pKT25lib$_{MS}$ library for transformation and the amount of transformation mixture were optimized to obtain approximately 300 colonies per plate and in total of 42 000 colonies. The plates were incubated for 48 hours at 30° and the transformants were screened for the blue colonies indicating the interactions of a protein fragment with the ParA protein. During the pre-selection 270 clones were re-streaked to confirm positive result. Next, plasmid DNA from 40 positive clones was isolated and presence of the insert in library clones was tested by PCR (with pKT25Fw i pKT25Rv primers). Obtained plasmid DNA was next used for retransformation with pUT18*parA* and to exclude false negatives, with pUT18C. The positive clones were sequenced.

Next, the whole *msmeg_5597* was cloned to pUT18C and pKT25 BTH vectors. For PCR amplification of *M. smegmatis papM* gene MSMEG_5597_XbaI_BamHI_Fw and MSMEG_5597_KpnI_EcoRI_Rv primers were used *M. smegmatis* mc$^2$ 155 chromosomal DNA as the template. The PCR product and vectors: pUT18C and pKT25 were cut by XbaI and KpnI restriction enzymes and ligated. The ligation mixtures were used for transformation of DH5α cells. The obtained clones were verified using PCR, enzyme digestion and sequencing.

To analyse the interaction between the studied proteins in the BTH system, pUT18C and pKT25 derivatives with analysed genes were transformed into *E. coli* BTH101 and plated on LB containing with ampicillin, kanamycin, X-gal and IPTG. After 2 days of incubation at 30°C, the selected representative transformants were re-streaked on LB containing X-gal, kanamycin, ampicillin, and IPTG.

**Constructs for *E. coli* colocalization**

pETDuet-$pknB_{KD}$ was constructed by amplifying the fragment of *M. smegmatis pknB* gene with PknB_Ms_Fw_HindIII and PknB_Ms_Rv_EcoRI primers and the *M. smegmatis* $mc^2$ 155 chromosomal DNA as the template. PCR product and pETDuet-1 vector were digested with HindIII and EcoRI restriction enzymes and ligated.

To obtain pETDuet-1 derivatives with *mcherry-divIVA* or $pknB_{KD}$ and *mcherry-divIVA*, the *ics-mcherry-divIVA* gene was PCR amplified with pJP108divIVA as a template and Cherry_Fw_Nde and Div_revSnaBI primers. The PCR product was digested with NdeI and SnaBI enzymes and ligated into pETDuet-1 or pETDuet-$pknB_{KD}$ vectors digested with NdeI and EcoRV, yielding pETDuet-*mcherry-divIVA* or pETDuet- $pknB_{KD}$ *mcherry-divIVA* respectively.

To construct pETDuet-*ics-mcherry, icsA-mcherry* gene was PCR amplified with pJP108*divIVA* as a template and Ics_Fw_NdeI and Cherry_rvSnaBI primers. The PCR product was digested with NdeI and SnaBI enzymes and ligated into pETDuet-1 vector digested with NdeI and EcoRV.

To construct pACYC-*papM-mT2*, first *papM* gene was amplified using 5597_Fw_NcoI and 5597_linker_Rv primers and the *M. smegmatis* $mc^2$ 155 chromosomal DNA as a template. *mTurquoise2* gene was amplified using mT2_Fw and mt2_Rv_EcoRI primers and p2Nil-*lsr2-mT2* as a template. To linearized the vector, the pACYC-Duet-1 vector was digested with NcoI and EcoRI restriction enzymes. The PCR products were assembled into the pACYC-Duet-1 vector using Gibson Assembly Kit (NEB) (xxx).

To construct pACYC derivatives containing *papM-mT2* and *egfp-parA*, the *egfp-parA* gene was excised from pACYC-*egfp-parA* vector with NdeI and PacI restriction enzymes and cloned into pACYC-*papM-mT2* vector digested with the same enzymes.

All the obtained constructs were analysed by PCR, restrictive digestion and sequencing.

**Preparation of *E. coli* constructs for protein overproduction and purification**

The pET28a*papM* construct was prepared by excision of the *papM* gene from the pKT25*papM* construct with BamHI and EcoRI enzymes and ligation into the pET28a*Φ* vector cut with the same enzymes. The ligation mixtures were used for transformation of DH5α cells. The obtained transformants were verified by PCR reaction using MSMEG_5597Fw and MSMEG_5597Rv primers and digestion with the SacI and BamHI restriction enzymes. The obtained, positive clones were transformed into BL21 (DE3) *E. coli* cells.

**PapM protein purification**

The *M. smegmatis* ParA protein was purified as an N-terminally GST-tagged protein using a GST-fusion purification system (GE Healthcare) as described earlier (Pióro 2022). MSMEG_5597 (PapM) protein was purified as a C-terminal His-tagged fusion protein using nickel-affinity chromatography. Briefly, cells from a 0.8 l culture were lysed by sonication in 50 ml of PBS supplemented with 10 mM imidazole. The obtained lysate was clarified by centrifugation (15 000 g, 15 min, 4 °C) and incubated (at 4 °C, overnight) with a 1 ml bed volume of Ni-NTA agarose resin (Qiagen). Next, the resin was washed three times with PBS supplemented with 50 mM imidazole, and bound proteins were eluted with PBS supplemented with 500 mM imidazole.

**Western blotting analysis**

For Western blot analysis was used 10 µg of lysate for each analyzed *E. coli* strain. Proteins were separated by SDS-PAGE electrophoresis. After electrophoresis, proteins were transferred to a nitrocellulose membrane (Millipore), which was blocked for 1 hour with 3% (wt/vol) BSA in TBST buffer (TBS with 0.05% Tween 20). The membrane was washed with TBST buffer, incubated with agitation for 1 hour at room temperature with anti-[p]threonine antibody rabbit monoclonal (diluted 1:500; Sigma-Aldrich), or anti His- antibody (diluted 1:1000; ThermoFisher Scientific) then washed three times with TBST, and incubated with agitation for 1 h at 20°C with goat anti-rabbit IgG conjugated to horseradish peroxidase (1:5000; Santa Cruz Biotechnology). The membrane was washed three times in TBST solution, the proteins were visualized with the SuperSignal West Pico Plus chemiluminescence substrate (ThermoFisher Scientific), and the results were captured using a ChemiDoc XRS system (Bio-Rad).

**Electophoretic mobility shift assay**

The whole plasmid pUC19 (2686 bp) was used for the EMSA (Electrophoretic Mobility Shift Assay) experiment. Protein binding to DNA was carried out in DNA binding buffer (50 mM Tris-HCl pH 7.5, 150 mM NaCl, 10 mM magnesium acetate, 0.02% Tween-20, 5 % [v/v] glycerol, 1mg/ml BSA). Increasing concentrations of His-PapM protein in the range of 0-2µM were incubated (20 minutes, 20°C) with 50 fmol of DNA in a final reaction volume of 20 µl. The samples were then analyzed electrophoretically. Electrophoresis was carried out in 0.8% native agarose gel in 1x TBE buffer at 4°C, overnight at a constant voltage of 4-5V/cm. To visualize the DNA, the gel was stained with ethidium bromide.

***M. smegmatis* strain construction**

All vectors for *M. smegmatis* transformations were prepared in *E. coli* DH5α. For construction of *M. smegmatis* mc$^2$ 155 strain with deletion of *papM* gene p2NilΔ*papM* was constructed. The sequence flanking 5' end of *papM* (847 bp) was amplified using Msmeg5597_F1_HindIII_Fw and Msmeg5597_F1_BamHI_Rv primers and cloned into the HindIII/BamHI sites of p2Nil vector to create p2Nil*papM*F1. Next, the region flanking downstream of *papM* 3' (907 bp) was amplified using Msmeg5597_F2_BamHI_Fw and Msmeg5597_F2_PacI_Rv primers and cloned into the BamHI/PacI sites of p2Nil*papM*F1, to yield p2Nil*ΔpapM*. Next, the pGOAL17 cassette was cloned into the PacI site of p2NilΔ*papM* to create p2NilΔ*papM*GOAL. The obtained vector was used to transform wild-type *M. smegmatis* mc$^2$ cells, yielding the IM01 (Δ*papM*) strain. To construct the strain with double deletion of *parA* and *papM* genes, p2NilΔ*papM*GOAL vector was used to transform *M. smegmatis* KG22 (Δ*parA*) cells, yielding the IM02 (Δ*papM* Δ*parA*) mutant strain. For the transformation of electrocompetent *M. smegmatis* cells, the plasmid DNA was treated with NaOH/EDTA (0.2 mM/0.2 mM). Transformants were plated on NB plates and selected for kanamycin resistance. The Kan$^R$ SCO (single-crossover recombinant) colonies were blue and sensitive to sucrose (2%). The SCO colonies were further plated on NB without selection. Cells were then resuspended in a liquid medium and serial dilutions were plated onto NB plates supplemented with sucrose and X-Gal. The selected double-crossover (DCO) mutants were white, Kan$^S$ and resistant to sucrose. PCR was used to distinguish between wild-type and DCO mutant cells.

For the construction of *M. smegmatis* complementation and *papM* overexpressing strains derivatives of the mycobacteriophage L5-based integration-proficient vector pMV306p$_{ami}$ were used. *M. smegmatis papM* gene was PCR amplified with the 5597_pami_XbaI_Fw and 5597_pami_KpnI_Rv primers and the *M. smegmatis* mc$^2$ 155 chromosomal DNA as a template. PCR products as well as pMV306p$_{ami}$*Ф* were digested with XbaI and KpnI restriction enzymes and ligated. The obtained construct was verified by PCR, restriction cloning and sequencing. Next, the obtained construct, as well as the control vector pMV306p$_{ami}$Ø, were used to transform wild-type, KG22 (Δ*parA*) (Ginda et al., 2013) and IM02 (Δ*papM* Δ*parA*) strains using the standard protocol, yielding IM12 (wt pMVp$_{ami}$*papM*), IM11 (Δ*parA*, pMVp$_{ami}$Ø), IM13 strains (Δ*parA*, pMVp$_{ami}$*papM*) and IMPW1 (Δ*papM* Δ*parA* pMVp$_{ami}$*papM*), respectively. Vectors were introduced into *M. smegmatis* competent cells by electroporation and transformants were selected on a medium containing kanamycin (50 µg/ml) and verified by PCR and western blotting.

Derivatives of strain IM02 (Δ*papM* Δ*parA*), producing EGFP-ParA or PAmCherry-ParA were constructed by the introduction of pMVp$_{nat}$*egfp-parA* plasmid (Ginda et, al., 2013) or pMVp$_{nat}$PA*mcherry-parA* (Pióro et al., 2019), yielding IM14 (Δ*papM* Δ*parA* pMVp$_{nat}$*egfp-parA*) or IM15 (Δ*papM* Δ*parA* pMV306p$_{nat}$*PAmcherry–parA*) strains, respectively. Obtained recombinants were checked by PCR, Western blotting and the production of fluorescent protein fusions was additionally verified by semi-native SDS-PAGE and subsequent detection with Azure 600 Imaging System (Azure Biosystems).

For the construction of the strains producing mCherry-DivIVA in different *M. smegmatis* genetic backgrounds, first, the *mcherry-divIVA* gene was cut out from pETDuet-*mcherry-divIVA* vector with NdeI and PacI restriction enzymes and cloned into pKW08p$_{tet}$*Φ* vector digested with the same enzymes. The resulting construct was analysed by PCR using pKW08F2 and pKW08RV2 primers, restrictive digestion and sequencing. Next, the obtained construct, pKW08p$_{tet}$*mcherry-divIVA*, was introduced into wild-type *M. smegmatis* mc$^2$ 155, IM01(Δ*papM*) and KG22 (Δ*parA*), yielding IM16 (wt pKW08p$_{tet}$*mcherry-divIVA*), IM17 (Δ*papM* pKW08p$_{tet}$*mcherry-divIVA*), IM18 (Δ*parA pKW08p$_{tet}$*mcherry-divIVA*) strains respectively. Mutants were analysed by PCR, DNA sequencing and/or Western blotting.

**RNA isolation, Reverse-Transcription and Quantitative PCR (RT-qPCR)**

For RT-qPCR reaction, RNA was isolated with Trizol LS reagent (Invitrogen) as described previously (Płociński et al., 2019). Briefly, 50 ml of *M. smegmatis* culture was centrifuged at 5,000×g for 10 min at 4°C and next the cells were resuspended in 300 µl of water and lysed by bead-beating with the MP FastPrep system (MP Biomedicals) using the program: 2 × 45 s, 6.0 m/s with 5 min intervals on ice. RNA was purified and treated with DNase I (RapidOut DNA Removal Kit, Invitrogen) according to the manufacturer's protocol. RNA quantity and integrity were checked by electrophoresis using an agarose gel.

A total of 500 ng of RNA was used for cDNA synthesis using the Maxima First Strand cDNA Synthesis Kit (Thermo Fisher Scientific) in a final volume of 20 µl. The original manufacturer protocol was modified for GC-rich *M. smegmatis* transcripts by extending the first-strand synthesis time up to 30 min and increasing the temperature for synthesis to 65°C. Subsequently, the resulting cDNA was diluted (1:5) and used for quantitative PCRs performed with PowerUp SYBR Green Master Mix (Applied Biosystems). The relative level of a particular transcript was quantified using the comparative ΔΔCt method and the *sig*A gene as the endogenous control (StepOne Plus Real-time PCR system, Applied Biosystems). The optimized oligonucleotides used in this study were synthetized by Genomed (Table S2).

**Homology modelling and analysis**

The homology models where predicted using the multimer mode in a standalone version of Alphafold (Evans et al., 2022). Structures were visualized in ChimeraX (Pettersen et al., 2021).
